# Effects of compassion training on brain responses to suffering others

**DOI:** 10.1101/616029

**Authors:** Yoni K. Ashar, Jessica R. Andrews-Hanna, Joan Halifax, Sona Dimidjian, Tor D. Wager

**Affiliations:** University of Colorado, Boulder, Department of Psychology and Neuroscience. Boulder, Colorado, USA; University of Arizona, Department of Psychology. Tucson, Arizona, USA; Upaya Institute and Zen Center. Santa Fe, New Mexico, USA

**Author notes:** Please address correspondence to: Tor D. Wager, Department of Psychology and Neuroscience, University of Colorado, Boulder, 345 UCB, Boulder, CO 80309.

**Keywords:** compassion training, empathy, mindfulness, placebo, burnout

## Abstract

What are the active ingredients and brain mechanisms of compassion training? To address these questions, we conducted a three-armed randomized trial (*N* = 57) of compassion meditation (CM). We compared a four-week CM program delivered by smartphone application to i) a placebo condition, in which participants inhaled sham oxytocin, which they were told would enhance compassion, and ii) a familiarity control condition, designed to control for increased familiarity with suffering others. Functional MRI was collected while participants listened to narratives describing suffering others at pre- and post-intervention. CM increased brain responses to suffering others in the medial orbitofrontal cortex (mOFC) relative to both the placebo and familiarity control conditions, and in the nucleus accumbens relative to the familiarity control condition. Results support the specific efficacy of CM beyond effects of expectancy, demand characteristics, and increased familiarity with suffering others, and implicate affective and motivational pathways as brain mechanisms of CM.

**Author Note:** Funded by the John Templeton Foundation’s Positive Neuroscience project (PIs Wager and Dimidjian), with additional support from NIH R01 R01DA035484 (PI Wager). Gratitude to research assistants Jenifer Mutari, Robin Kay, Scott Meyers, Nicholas Peterson, and Brandin Williams for help with data collection.

---

Compassion is a vital societal and interpersonal resource facilitating caring and cooperative behavior (Goetz, Keltner, & Simon-Thomas, 2010), and it is widely recognized a virtue. Recently, scientific interest has turned toward the cultivation of compassion. Evidence is accumulating that training programs can enhance compassion in community samples (Galante, Galante, Bekkers, & Gallacher, 2014), medical providers (van Berkhout & Malouff, 2015), and patient populations (Gilbert, 2014). Many of these compassion training programs have been based on compassion meditation (CM), a secular meditation practice with Buddhist origins. In the scientific literature, CM has been viewed as a practice that enhances compassion by influencing appraisals of and emotional responses to suffering others (Ashar, Andrews-Hanna, Yarkoni, et al., 2016; Dahl, Lutz, & Davidson, 2015, 2016; Engen & Singer, 2015a; Weng, Schuyler, & Davidson, 2017).

Here, we focus on two open questions regarding CM. First, its “active ingredients” remain unclear: the observed effects of CM could be due to expectations of increased compassion, demand characteristics, and/or other “non-specific” components. Although some studies report specific effects of CM relative to other meditation practices (e.g., Kok & Singer, 2017; Valk et al., 2017) or to empathy training (Klimecki, Leiberg, Ricard, & Singer, 2014), meta-analyses have found relatively weak support for the specific efficacy of CM in comparison to active controls (Galante et al., 2014; Kreplin, Farias, & Brazil, 2018). More comparisons of CM against control groups matched on non-specific factors, such as placebo conditions, are needed.

A second question pertains to the neurobiological mechanisms of CM. Previous brain imaging studies have found that, relative to control conditions, CM has led to increased brain activity in two different sets of brain regions: 1) the mesolimbic dopaminergic system, which includes the ventral striatum, ventral tegmental area (VTA), medial orbitofrontal cortex (mOFC), and ventromedial prefrontal cortex (vmPFC), and is closely tied to reward and motivation processes (Engen & Singer, 2015b; Klimecki et al., 2014; Singer & Klimecki, 2014), and 2) regions linked to mentalizing and reappraisal, including the dorsomedial prefrontal cortex (dmPFC), dorsolateral prefrontal cortex, and the inferior parietal lobule (Ashar, Andrews-Hanna, Dimidjian, & Wager, 2016; Mascaro, Rilling, Tenzin Negi, & Raison, 2013a; Weng et al., 2013). The effects observed in these distinct sets of brain regions parallel a current debate in the meditation literature: whether CM primarily affects more “affective” or more “cognitive” processes (Dahl et al., 2015, 2016; Engen & Singer, 2015a). More neuroimaging studies are needed to advance understanding of the psychological and neurobiological mechanisms of CM.

To address these questions, we conducted a three-armed randomized controlled trial of CM, with each group receiving a four-week intervention delivered daily by a mobile iPod application. The CM program aimed to teach participants skills for staying engaged with others’ suffering without becoming emotionally overwhelmed, as research suggests that feeling compassion is experienced as effortful and is often avoided (Cameron, 2017; Cameron et al., 2019; Zaki, 2014). During the guided meditations, participants listened to a true story describing a suffering person, looked at a corresponding face photograph, and engaged in particular CM techniques directed toward the suffering person. A comparison group received a placebo intervention, to test for specific effects of CM beyond expectancy effects and demand characteristics. Participants in this group listened to the same stories daily while viewing corresponding face photographs after inhaling a nasal spray described to them as oxytocin, a compassion-enhancing drug – though it was actually saline. Additionally, a second comparison group simply listened to the same stories daily: we hypothesized that increased familiarity with suffering others alone could enhance compassion (“Familiarity” group). To investigate the neurobiological effects of CM, we collected functional MRI (fMRI) pre- and post-intervention. During scanning, participants listened to the same stories describing suffering others while viewing their associated face photographs.

## Method

### Participants

Out of 311 participants screened for eligibility, 71 healthy adults completed the baseline assessment between January and September of 2012. To be eligible, participants were required to self-report no history of major psychiatric illness, current mental health conditions, or breast-feeding (to maintain the oxytocin placebo deception). Standard fMRI exclusion criteria were applied (e.g., no metal in body, no claustrophobia). Participants also were required to have no previous experience with CM or Loving-Kindness Meditation and at least moderate interest in meditation, as we sought to investigate the effects of CM among healthy, interested novices (e.g., see also Segal et al., 2010; Williams et al., 2014). We also excluded participants who reported in advance that they would not donate any participation earnings to charities to ensure variability on a primary outcome of charitable donation. Thirteen participants who completed the baseline assessment were not eligible for randomization for a variety of technical reasons (see Figure S3 in Ashar et al., 2016), primarily excessive head motion during the baseline fMRI scan. Thus, *N* = 58 participants were randomized using a computer-generated randomization list, stratified by sex. One participant refused randomization to the placebo oxytocin condition due to unwillingness to use a nasal spray. Thus, the final sample included *N* = 57 analyzed participants, including *N* = 36 females, with M_age_ = 29.11 years, SD _age_ = 6.35 years, M_Subjective SES_ = 6.25 out of 10 and SD _Subjective SES_ = 1.65. Subjective SES was measured with a single-item measure (Adler, Epel, Castellazzo, & Ickovics, 2000). Additionally, one participant misunderstood the donation task instructions and was dropped from analyses of the charitable donation outcomes. Experimenters were blind to the participants’ assigned intervention for the pre-intervention assessment but not for the post-intervention assessment. Participant demographics and baseline characteristics are provided for each intervention condition in Table 1 of Ashar et al., 2016. Intervention effects on behavioral outcomes were reported in Ashar et al. 2016, and baseline only brain data in Ashar et al. 2017; the current manuscript reports intervention effects on brain activity, which have not previously been reported. Sample size was dictated by the available research funds.

Participants were compensated $100 for each MRI session, and an additional $1 for each day of the intervention that they completed. After completion of the study, OxyPla participants completed a questionnaire assessing the strength of their belief that they were actually taking oxytocin and were then debriefed regarding the nature of the deception and its purpose. The University of Colorado Institutional Review Board approved all procedures, including informed consent. No serious adverse events resulted from any of the intervention conditions.

### Materials and Procedures

#### Brain responses to suffering others

During fMRI scanning conducted before and after the four-week intervention, participants listened to 24 randomly ordered biographies describing true stories of suffering people, such as orphaned children, adults with cancer, and homeless veterans. Biographies were created from factual information posted on charity websites and recorded by one member of the research team as audio segments 26 to 33.5 s in duration. An authentic facial photograph of each person, also drawn from the charity website, was displayed alongside the audio biography. The people described in the biographies were balanced on age (child or adult), race (Black or White), and sex. Real stories and photographs were used to increase ecological validity. An example biography is: “*Jessica’s father abandoned his family and her mother was unable to support them alone, so they had to move into a homeless shelter. The shelter provided Jessica's mother with professional training and childcare. Eventually, the family moved into subsidized housing. Jessica and her sisters have been tremendously supportive of each other. Jessica has managed to stay in school and will finish the year with her class.*” To hear this biography while viewing a sample face photograph, as presented to our subjects, visit https://canlabweb.colorado.edu/files/jessica.mp4 (image copyright CC BY-SA 2.0). The text of all biographies is listed in Table S1 of Ashar et al. 2017, and audio and video recordings of all biographies are available for download at https://github.com/canlab/Paradigms_Public/tree/master/2016_Ashar_Empathy_CompassionMeditation. After each biography, participants provided ratings of empathic care or empathic distress (data not presented here).

Participants then heard abbreviated “reminder” biographies (8 - 11 sec) during fMRI scanning while viewing the face photograph of that person. An example reminder is: “*Jessica’s father abandoned his family. Her mother and sisters moved into a homeless shelter, which provided job training and childcare. Jessica will finish school this year with her class.*” Reminders were provided because pilot studies showed that participants had difficulty distinctly recalling each of the 24 biographies when asked to make donation decisions. Following each reminder, participants were given an option to donate a portion of their own experimental earnings to a charity helping that person, from $0 to $100 in $1 increments, as a measure of compassionate behavior. Between trials, participants were asked to press a button indicating which direction an arrow was pointing (left or right); this served as a non-social baseline comparison task. The duration of this inter-trial baseline task was jittered across trials, from 3 to 9 seconds. During the task, participants were asked to simply listen to the biographies, and CM participants at the post-intervention assessment were asked not to engage in CM while listening to the biographies, for greater comparability across conditions.

This task was completed over three fMRI runs of listening to biographies and rating feelings, followed by two runs of listening to biography reminders and making donations. The task is publicly available at https://github.com/canlab/Paradigms_Public/tree/master/2016_Ashar_Empathy_CompassionMeditation.

#### Self-reported and behavioral measures of compassion

Primary self-reported and behavioral outcomes were charitable donations, as described in the fMRI task above, and compassion-related feelings and attributions. Participants listened to the biographies a second time after scanning and provided ratings of feelings, attributions, and aspects of perceived similarity for each biography on a visual analog scale. These ratings included attributions of blame-worthiness for one’s suffering, attributions of how much a person would be benefitted by efforts to help them, and feelings of distress and tenderness. In prior work (Study 1 in Ashar, Andrews-Hanna, Yarkoni, et al., 2016), we reported that a linear combination of these feelings and attributions, termed Feeling-Attribution-Similarity (or “FAS”) scores, was strongly predictive of charitable donation. We applied this model to the data collected in this study, to generate FAS scores for each participant at pre- and post-intervention.

#### Interventions

After the baseline session, participants were randomized to one of the three interventions – Familiarity, Placebo oxytocin (OxyPla), or Compassion Meditation (CM) training – with *N*_CM_ = 21, *N*_OxyPla_ = 18, and *N*_familiarity_ = 18. The three interventions were delivered via iPod Touch applications developed by the study team and matched across conditions on structure and style. All participants were asked to complete a daily task for four weeks on the iPod Touch provided to them. A member of the study team placed three phone calls to participants during the intervention to address any concerns, ask about side effects in the OxyPla condition, and encourage compliance.

Participants in all three conditions listened to a biography of a suffering person every day while viewing a photograph of that person. Out of the 24 total biographies presented during the fMRI sessions, each participant listened to and viewed a set of 12 biographies across the four-week intervention period. The set of biographies presented to each participant during the intervention was randomly assigned and balanced across groups.

#### Compassion Meditation

The CM program was designed to enhance both compassion for suffering others as well as equanimity. The emphasis on equanimity aimed to help prevent emotional overwhelm from others’ suffering, thereby promoting sustainable compassionate responding (Halifax, 2012). A theme of the meditation recordings was “*soft front, strong back*,” which served as both a metaphor and an embodied approach to being sensitive to and engaged with others’ suffering while remaining emotionally grounded. Meditations included a focus on grounding in the body and connection with the earth as a foundation, perspective taking practices, visual imagery (e.g., imagine the suffering person as a small child), and repetition of compassion-related phrases directed toward the suffering person (e.g. “may you find peace”). The meditations also asked participants to direct these practices toward themselves, to enhance self-compassion. Participants were asked to practice meditation for about 20 minutes daily and were provided with a new guided meditation practice at the start of each of the four weeks. At a specified point during each daily meditation, participants would hear one of the biographies described in the fMRI task above and were asked to meditate on that person specifically. A more detailed description of the CM program is provided in Supplemental Material of Ashar et al. 2016.

#### Placebo oxytocin intervention

The placebo oxytocin (OxyPla) intervention was designed to control for placebo effects related to CM, such as a) participant expectations of increased compassion naturally created by CM, and b) demand characteristics created by completing a CM intervention in a research context (e.g., wanting to satisfy perceived researcher objectives by exhibiting increased compassion). Prior to listening to each daily biography, participants were instructed to inhale a nasal spray labeled as oxytocin. Participants were also provided with scientific information sheets describing oxytocin’s ability to enhance compassion.

#### Familiarity intervention

Participants in the Familiarity condition simply listened to one biography of a suffering person daily. This condition was designed to control for the increased familiarity with suffering others inherent in the CM practice, as familiarity with suffering people could increase liking and enhance compassion (Zajonc, 2001).

#### Daily attention-to-task check

After each daily intervention task, participants in all conditions responded to a multiple-choice question designed to test whether they had adequately paid attention to the biography. Participants were asked to indicate the primary hardship afflicting the individual described in the biography they had heard that day (e.g., *What was Robert’s primary hardship? A) AIDS, B) Cancer, or C) Homelessness*). Participants also provided ratings of mood each day (data not presented here).

### Analyses

#### fMRI data acquisition and preprocessing

Images were acquired with a 3.0 T Siemens Trio Tim magnetic resonance imaging scanner using a 12-channel head coil. Twenty-six 3.0-mm-thick slices (in-plane resolution 3.4 × 3.4 × 3.0, 1 mm gap, ascending sequential acquisition) extended axially from the mid-pons to the top of the brain, providing whole-brain coverage (TR = 1.3 s, TE = 25 ms, flip= 75°, field of view = 220 mm, matrix size = 64 × 64 × 26). High-resolution structural scans were acquired prior to the functional runs with a T1-weighted MP RAGE pulse sequence (TR = 2530 ms, TE = 1.64 ms, flip = 7°, 192 slices, 1 × 1 × 1 mm). Parallel image reconstruction (GRAPPA) with an acceleration factor of 2 was used.

Before fMRI preprocessing, volumes were identified as outliers on signal intensity using Mahalanobis distances (3 standard deviations) and dummy regressors were included as nuisance covariates in the first level models. Functional images were corrected for differences in the acquisition timing of each slice and were motion-corrected (realigned) using SPM8. Twenty-four head motion covariates per run were entered into each first level model (displacement in six dimensions, displacement squared, derivatives of displacement, and derivatives squared). Structural T1-weighted images were then coregistered to the mean functional images using SPM8’s iterative mutual information-based algorithm. Coregistered, high-resolution structural images were warped to Montreal Neurologic Institute space (avg152T1.nii); these warping parameters were applied to the functional data, normalizing it to MNI space, and interpolated to 2×2×2 mm^3^ voxels. Lastly, functional images were smoothed with an 8 mm FWHM Gaussian kernel. A 220-second high-pass filter was applied during first-level analysis.

#### fMRI analyses

Our analyses focused on the period of listening to biography reminders, which were briefer (11.5 seconds) than the initial biography listening period (33.5 seconds) and more proximal to charitable donation decisions. We estimated a general linear model (GLM) using SPM8 for each participant including the nuisance covariates generated in preprocessing and two regressors of interest: listening to biography reminders (11.5 sec), and the charitable donation decision period (5 sec), each convolved with the standard hemodynamic response function. The jittered-duration inter-trial interval served as the model intercept. We then computed contrast images for the [listen – baseline] comparison for every subject at pre- and post-intervention. We subtracted the pre-intervention image from the post-intervention image to estimate pre-to-post-intervention changes in brain responses to stories of suffering. These images were used for both the ROI and whole-brain analyses described below.

We tested for CM vs. Familiarity and CM vs. OxyPla group differences. These two contrasts estimated the specific effects of CM relative to a condition simply repeatedly exposing participants to others’ suffering, and the effects of CM relative to a placebo condition. For archival purposes, we also estimated the OxyPla vs. Familiarity comparison to characterize placebo effects on brain activity; these results are reported in the supplementary material (Supplementary Table 2, Figure S6).

#### Region of interest analyses

Previous CM studies have reported effects primarily in the mesolimbic dopaminergic pathway or in mentalizing-related regions (reviewed above). We tested these *a priori* neuroanatomical hypotheses in several key regions of interest (ROIs) identified by previous studies: the VTA, nucleus accumbens (NAcc), mOFC, vmPFC, dmPFC, and temporoparietal junction (TPJ). We used one-tailed tests, given the *a priori* directional hypotheses afforded by prior literature that CM would lead to increased activity relative to controls. Masks for the VTA and NAcc were taken from a high-resolution subcortical atlas (Pauli, Nili, & Tyszka, 2018) and masks for the vmPFC, dmPFC, TPJ, and mOFC were adopted from a recent multi-modal cortical parcellation (regions 10, 9, ‘PFm’, and ‘OFC’ in Glasser et al., 2016, respectively). Given our small sample size, limited statistical power, and *a priori* hypotheses, we did not correct for multiple comparisons across ROIs.

#### Whole brain analyses

To further investigate intervention effects beyond the regions specified in the ROI analyses, we conducted a whole-brain robust regression (Wager, Keller, Lacey, & Jonides, 2005) estimating group differences in pre-to-post-intervention changes in brain activity. We also conducted a robust regression estimating average pre-to-post-intervention changes *within* each group. This characterized absolute pre-to-post-intervention increases or decreases in brain responses to suffering others, for each group independently.

Whole-brain analyses were subjected to a voxelwise threshold of *p* < .001 uncorrected with a cluster-defining threshold of 10 voxels – a commonly used threshold (Woo, Krishnan, & Wager, 2014) providing a balance between type I and type II error rates (Lieberman 2009). No voxels survived false discovery rate (FDR) correction at *q* < .05. Future studies with larger sample sizes will allow for more stringent statistical thresholding. All analyses were conducted within a gray matter mask. Neuroanatomical labeling was conducted using the freely available CanlabCore tools, which pools anatomical labels from across number of published atlases (see https://canlabcore.readthedocs.io/en/latest/moduleslist.html#@region.table), and with reference to the Harvard-Oxford cortical atlas as well. Contrast images for each subject at each time point, behavioral outcomes, and code for all analyses are publicly available here: https://github.com/yonestar/effects_of_CM_on_brain.

#### Intervention effects on brain models of empathic care and distress

We also tested the effect of the interventions on previously published brain models of empathic care and distress (Ashar, Andrews-Hanna, Dimidjian, & Wager, 2017), which served as an *a priori* brain measure of emotional responses to suffering others. We hypothesized that the CM group would exhibit pre-to-post-intervention increases in these brain models, relative to control conditions. We computed the cosine similarity between the care and distress models and each subjects’ contrast images at both pre-and-post-intervention. We then computed a pre-to-post-intervention change score for each participant for both the empathic care and empathic distress models. We submitted these change scores to a two-sample *T*-test testing for CM vs. OxyPla and CM vs. Familiarity group differences.

## Results

### Behavioral results

We first summarize the previously published (Ashar et al., 2016) behavioral outcomes from the trial, to provide a context for the fMRI outcomes that are the focus of this manuscript.

#### Participant compliance and expectations

Intervention compliance, as logged by the intervention iPod applications, was high across groups, *M* = 24.16 days completed out of 28 possible days. There were significant group differences in compliance, *F*(2, 54) = 16.70, *p* < .001, which were driven by lower completion rates among CM participants, *M*_CM_ = 20.76 days, 95% CI = [18.81, 22.71], perhaps due to the increased time and/or cognitive-emotional demands of CM. Similarly, performance on the daily attention-to-task questions was near ceiling across groups, *M* = 98% correct, although there were significant group differences in correct responding, *F*(2, 54) = 6.00, *p* = .004, which were driven by slightly lower performance in the CM group, *M*_CM_ = 95% correct, 95% CI = [0.93, 0.98]. Overall, these data indicate that rates of intervention compliance and attention were generally high across groups, notwithstanding the fact that the interventions were delivered by smartphone applications. The high rates of compliance may have been due to the monetary incentive provided for completing the intervention, increased participant engagement due to the brain imaging component of the study, or other study features.

Pre-intervention expectations of increased compassion were moderately high across groups, *M* = 5.00 on a 0 to 10 scale, 95% CI [4.40, 5.60]. There were significant group differences between the three groups, *F*(2, 49) = 6.06, *p* = .004. This was driven by lower expectations in the Familiarity condition, *M* = 3.69 out of 10, 95% CI [2.49, 4.88]. Expectations in the Familiarity condition were statistically lower relative to the CM condition, *T*(32) = 3.28, *p* = 0.003, 95% CI [0.86, 3.66] and relative to the OxyPla condition, *T*(31) = 1.97, *p* = .06, 95% CI [−0.05, 2.91]. Direct comparison of expectations in the CM condition and the OxyPla condition showed no statistical difference, *T*(33) = 1.32, *p* = .19. Overall, these results suggest that CM does generate expectations of increased compassion, and that the placebo manipulation also succeeded in generated expectations of increased compassion.

#### Changes in compassion and donation

Relative to Familiarity participants, CM participants increased in both FAS scores and charitable donations from pre- to post-intervention, Hedge’s *g*_FAS_ = 1.17, 95% CI [0.69, 1.76], and g_donation_ = 0.89, 95% CI [0.35, 1.47], *M*_donation difference_ = $8.17, 95% CI [$2.22, $14.11]. Relative to OxyPla participants, CM participants significantly increased in FAS scores, *g*_FAS_ = 0.69, 95% CI [0.10, 1.34], but the difference in donations was not statistically significant, g_donation_ = 0.48, 95% CI [−0.14, 0.97], *M*_donation difference_ = $5.95, 95% CI [$−2.33, $14.23]. Examining patterns of absolute pre-to-post-intervention change within group, we found that CM participants increased in FAS scores, *g* = 0.61, 95% CI [0.27, 1.03], and did not statistically change in donation amounts, *g* = 0.19, 95% CI [-0.27, 0.60]. OxyPla participants did not statistically change in either outcome, *g*_FAS_ = −0.03, 95% CI [−0.56, 0.41], *g*_donation_ = −0.27, 95% CI [-0.58, 0.22]. Familiarity participants decreased in both outcomes, *g*_FAS_ = −0.53, 95% CI [−0.95, −0.18], g_donation_ = −0.67, 95% CI [−1.06, −0.36] (Figure 2c). Overall, this profile of behavioral results suggests that: a) Familiarity participants decreased in compassion over time, b) CM buffered against this decrease and led to increased compassion, and c) the effects of CM were partly but not fully attributable to placebo.

### fMRI Results

#### ROI analyses

We tested for group differences in two sets of ROIs that have been previously implicated in CM neuroimaging studies: ROIs along the mesolimbic dopaminergic pathway and ROIs linked to mentalizing processes. For the CM vs. OxyPla comparison, no significant differences were observed. For the CM vs. Familiarity comparison, significant group differences were observed in the mOFC and the NAc. The ROIs and results are depicted in Figure 1 and Table 1.

**Figure 1.**
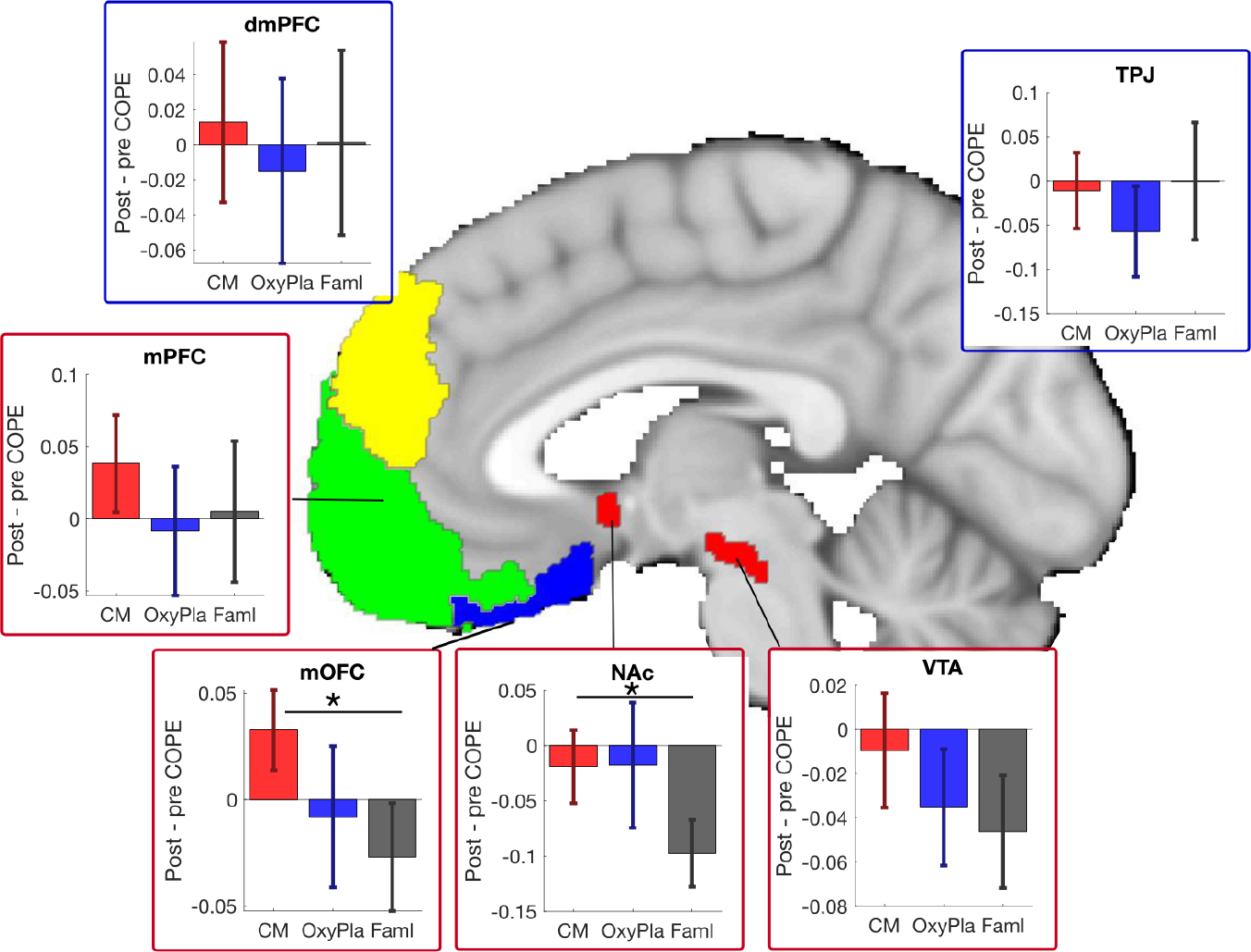
Pre-to-post-intervention changes in brain responses to suffering other in regions previously implicated in CM research, including regions along the mesolimbic dopaminergic pathway (red outlines) and regions linked to mentalizing processes (blue outlines). Subplots depict contrasts of parameter estimates (COPEs) for pre-to-post-intervention changes for each of the three groups: compassion meditation (CM), placebo oxytocin (OxyPla), and Familiarity (Faml). Error bars depict standard error; * = *p* < .05 one-tailed, given clear *a priori* directional hypotheses.

**Table 1.**
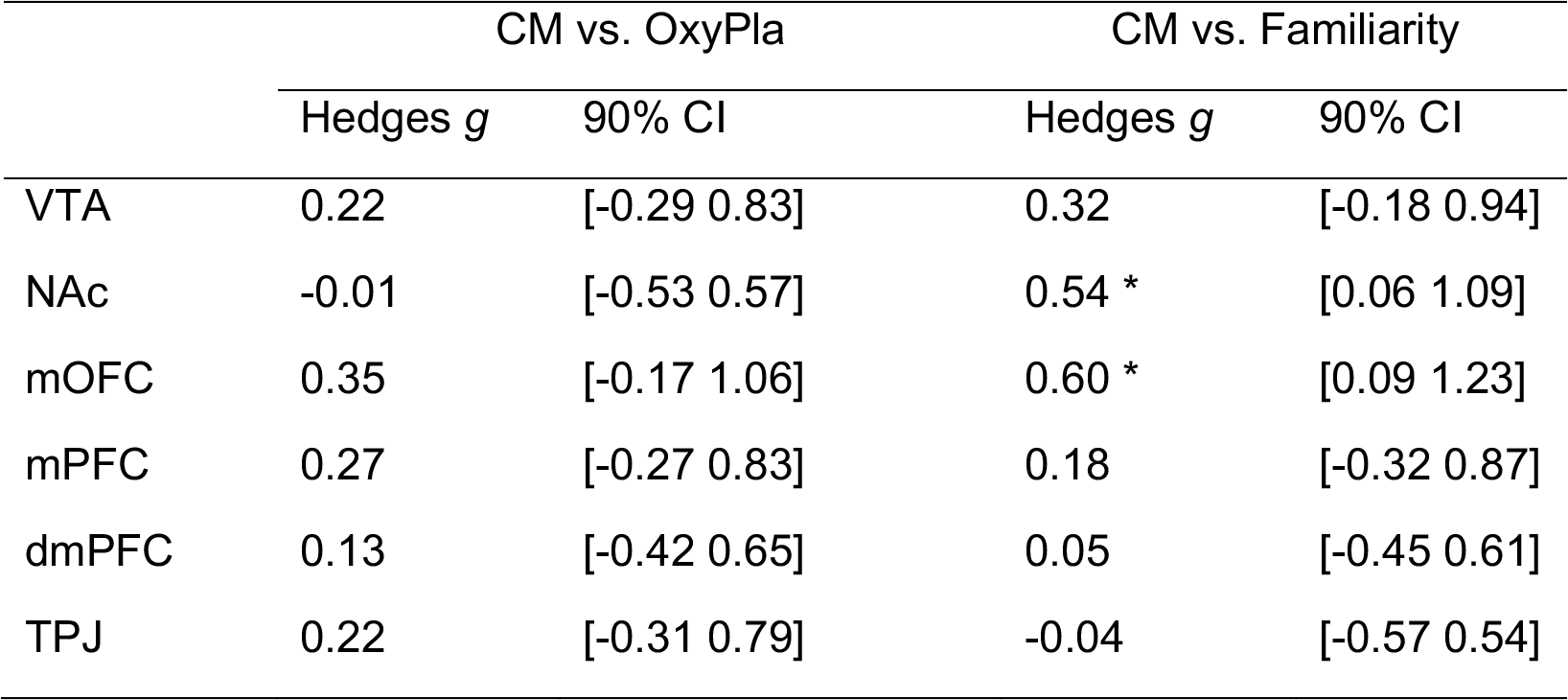
Results from ROI analyses, corresponding to Figure 1. * = *p* < .05 one-tailed, given clear *a priori* directional hypotheses.

#### Whole brain analyses

We compared pre-to-post intervention changes in brain responses to stories of suffering for CM vs. two control conditions (Figure 2a). The CM vs. OxyPla comparison yielded three clusters with relatively increased activity in the CM condition: the right mOFC and two occipital areas. The CM vs. Familiarity comparison yielded three clusters with relatively increased activity for the CM condition: the right mOFC, left superior temporal sulcus, and left parahippocampal cortex. Coordinates corresponding to brain regions showing significant effects are listed in Supplementary Table 1.

To better characterize intervention effects, we also examined pre-to-post-intervention change within each group (Figure 2b). CM participants exhibited increased brain responses to suffering others in the mOFC, mPFC, midbrain, and the right cerebellum. OxyPla participants exhibited no significant increases in brain activity over time, and some decreases in V3 and lateral prefrontal areas. Familiarity participants exhibited decreases across several prefrontal and subcortical structures, including: mOFC, insula, superior temporal sulcus, amygdala, hippocampus, hypothalamus, putamen, and globus pallidus. Coordinates corresponding to brain regions showing significant effects are listed in Supplementary Table 2.

Three-dimensional interactive images of all results are available at https://neurovault.org/collections/4766/ and figures showing full-brain results are provided in Figures S1 – S6.

**Figure 2.**
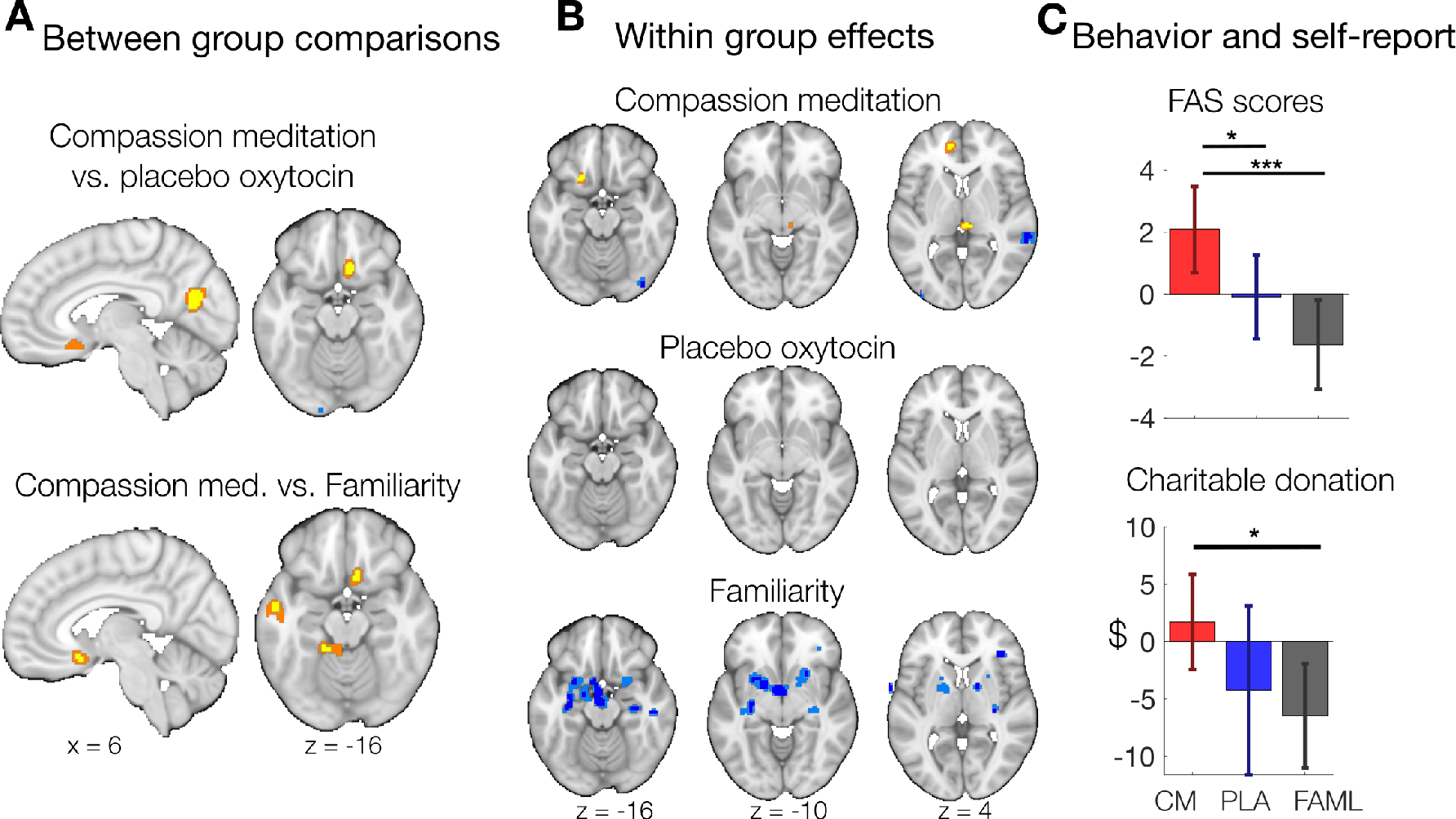
Pre-to-post-intervention changes in whole-brain and behavioral responses to suffering others. **A)** Comparisons of CM to control conditions for pre-to-post intervention changes in brain responses to audio-video narratives describing suffering others. **B)** Absolute pre-to-post-intervention changes within each condition. Yellow/orange areas showed increases from pre- to post-intervention, and blue areas showed decreases. Voxels meeting a threshold of *p* <.001 uncorrected are shown in yellow and dark blue, respectively. Adjacent voxels meeting a threshold of *p* < .005 are shown in orange/light blue, for visualization purposes. **C)** Pre-to-post-intervention changes in charitable donations and in a composite index of compassion-related feelings and attributions (“FAS scores”, see main text). Error bars show 95% confidence intervals; * = *p* < .05, ** = *p* < .01, *** = *p* < .001.

#### Test of buffering hypothesis

Results from the within-group and ROI analyses indicated decreased brain responses in the Familiarity group across a range of brain structures. These decreases not observed in the CM or OxyPla groups, suggesting that these interventions may have “buffered” against the decreases observed in the Familiarity group. We conducted a follow-up analysis to directly test the hypothesis that the Familiarity group exhibited a neuroanatomically reliable decrease, which was buffered against by the CM and/or OxyPla interventions. We conducted a cross-validated analysis selecting the most decreased voxels within a training subset of Familiarity participants. We then compared pre-to-post-intervention changes in those voxels in held-out data across the three groups. Results were compared to a null distribution created by permuting the group labels and repeating the analysis 10,000 times. We found a small, non-significant “buffering” effect of CM vs. Familiarity in the expected direction, *g* = .22, *p* > .4.

#### Effects on *a priori* brain models of empathic care and distress

Effects of the intervention were in the expected direction, with CM participants exhibiting increases in the neural signature response (i.e. pattern expression) for both empathic care and empathic distress relative to both OxyPla and Familiarity participants. However, the group by time interactions were not statistically significant (*p*s ranging from .12 - .72, for the CM vs. Familiarity comparison and the CM vs. OxyPla comparison, for the empathic care and distress patterns).

## Discussion

CM has received recent scientific interest as a promising intervention for enhancing compassion. Our present investigation of CM centered around two questions: What are the effects of CM on brain function? And what is the specific efficacy of CM, relative to conditions controlling for non-specific factors?

Our findings point to the mOFC as a region centrally affected by the CM intervention. ROI analyses identified a significant CM vs. Familiarity effect in the mOFC. Further, in whole brain analyses, the right mOFC was the only region to survive comparisons of CM to both control conditions, and a nearby region in the left mOFC demonstrated absolute pre-to-post-intervention increases in the CM group considered independently. Much research with human neuroimaging and neuronal recordings has strongly linked the mOFC to affective process such as reward and motivation (Haber & Knutson, 2010), empathic care for suffering others (Ashar et al., 2017), altruistic giving (Cutler & Campbell-meiklejohn, 2019), and social reward (Watson & Platt, 2012). This suggests that the changes we observed in this region may be related to the increased self-reported compassion and charitable donations observed in our participants.

Our mOFC results are in line with a number of previous CM findings. Engen and Singer (2015b) found that practicing compassion meditation during viewing of emotional videos activates the mOFC, as compared to practicing cognitive reappraisal strategies. Randomized trials comparing CM to empathy and memory trainings have also found that CM increased activity in mOFC areas overlapping or adjacent to the mOFC cluster reported here (Klimecki, Leiberg, Lamm, & Singer, 2012; Klimecki et al., 2014). Further, long-term meditators more strongly engage the mOFC when practicing CM relative to novice practitioners (Engen, Bernhardt, Skottnik, Ricard, & Singer, 2018). And yet, since these findings have all been reported by one research group, to our knowledge the current report is the first corroboration by an independent research group that CM leads to increased activity in the mOFC.

Our results bear on a recent debate regarding the mechanisms of CM: Do CM and related practices primarily induce changes in cognitive or affective processes (Dahl et al., 2015, 2016; Engen & Singer, 2015a)? Some data point to changes in mentalizing-related regions, such as the dorsomedial PFC (Mascaro, Rilling, Tenzin Negi, & Raison, 2013b) and the temporoparietal junction (Weng et al., 2013), while other studies report changes in the mesolimbic dopaminergic system including the ventral tegmental area, ventral striatum, and mOFC (Engen et al., 2018; Engen & Singer, 2015b; Klimecki et al., 2012, 2014), which have been linked to affect and motivation. In ROI analyses, we found significant CM vs. Familiarity group differences in the mOFC and the NAc, but not in the two mentalizing-related regions we tested. These supports the view that CM induces changes in affective and motivational processes. At the same time, cognitive and affective processes interact in empathy (Ashar, Andrews-Hanna, Dimidjian, et al., 2016; Kanske, Bockler, Trautwein, Lesemann, & Singer, 2016), and, our results cannot disambiguate between processes that are activated during the *practice* of CM vs. the *outcome* of CM (Dahl et al., 2016). Further, since CM represents a family of meditation practices, there may be substantial variability among different CM implementations (e.g., Cognitively-Based Compassion Training (Pace et al., 2009) may preferentially impact more cognitive processes).

What is the specific efficacy of CM, above and beyond placebo? Our results support the notion that the meditation practices included in the CM program had a specific effect: on a whole-brain level, we observed significant effects of CM on brain function relative to placebo. At the same time, in ROI analyses no significant CM vs. placebo differences were detected, while differences with the Familiarity condition were detected. These findings suggest that placebo effects account for some, but not all, of the effects of CM. This is in line with previous reports of the specific efficacy of CM in enhancing compassion as compared with other meditation practices (Kok & Singer, 2017; Valk et al., 2017) or empathy training (Klimecki et al., 2014). It also accords with reports of the specific effects of mindfulness meditation on brain function relative to sham meditation (Zeidan et al., 2015; Zeidan, Johnson, Gordon, & Goolkasian, 2010). Our findings also show that simply increasing familiarity with suffering others is not a key ingredient of CM, as CM separated from the Familiarity condition in several analyses.

A critical challenge is developing well-matched control conditions for meditation-based interventions (MacCoon et al., 2012). The comparison conditions included here did not control for some non-specific factors, such as time spent engaged in the intervention. And factors that we did attempt to control for, such as expectations of enhanced compassion, may not have been perfectly matched across groups (Boot, Simons, Stothart, & Stutts, 2013). Future studies including different control groups can continue to refine our understanding of the “active ingredients” of CM, potentially leading to the development of more effective compassion training interventions.

We also observed marked decreases in Familiarity participants across a range of prefrontal and subcortical regions, including the mOFC, amygdala, hippocampus, insula, and more. These decreases were not observed in the CM and/or placebo groups, suggesting a potential “buffering effect” of these interventions. However, inferences based on changes observed in one group but not the other are invalid, as the *interaction* must be significant (Nieuwenhuis, Forstmann, & Wagenmakers, 2011). In most of these brain structures, we did not detect a significant group by time interaction – perhaps because of our limited sample size.

The observed decreases in the Familiarity condition suggest a direction for future research: Can CM and/or placebo interventions can buffer against a decrease in compassion following repeated exposure to suffering? This idea has been raised by several authors as one with particular societal relevance (Ashar, Andrews-Hanna, Dimidjian, et al., 2016; Klimecki & Singer, 2011). For example, professional burnout among caregivers, such as doctors and nurses, poses a major societal problem, with about 50% of American physicians reporting at least one burnout symptom (Maslach & Leiter, 2016; Shanafelt, Goh, & Sinsky, 2017). One reason for caregiver burnout is the detachment, depersonalization and the erosion of meaning caused in part by repeated exposure to suffering others without tools for handling the accompanying emotional load (Maslach & Leiter, 2016). Our results suggest that CM could be investigated as a technique for maintaining positive emotional connections with patients, potentially reducing burnout (Mascaro et al., 2018) and enhancing patient outcomes (Kaptchuk et al., 2008; Rakel et al., 2011).

Similarly, compassion is also often reduced for multiple victims relative to a single victim (Cameron, 2017). This phenomenon poses a problem: it becomes difficult to motivate action addressing issues afflicting large numbers of suffering people (e.g., millions of starving children), while circumstances afflicting a single person (e.g., a single starving child) more easily garner compassion and helping behavior (Bloom, 2016). While many factors contribute to this “collapse of compassion” for multiple victims, experimental evidence suggests that fear of the costs of feeling compassion, both psychological and material, plays a central role (Cameron, 2017). Seen in this light, “compassion collapse” is a motivated choice to avoid feeling compassion (Zaki, 2014). Compassion training programs that teach skills for staying engaged with others’ suffering without becoming overwhelmed or overly distressed might reduce the fear of feeling compassion and help sustain compassion in difficult situations. Future studies directly investigating these questions are needed.

## Author contributions

T.D.W and S.D. developed the study concept. All authors contributed to study design. Testing and data collection were performed by Y.K.A. and research assistants under the supervision of Y.K.A. Data analysis and interpretation was conducted by Y.K.A. under the supervision of T.D.W. Y.K.A drafted the manuscript. All authors provided critical revisions and approved the final version of the manuscript for submission.

## References

Adler, N., Epel, E., Castellazzo, G., & Ickovics, J. (2000). Relationship of subjective and objective social status with psychological and physiological functioning: Preliminary data in healthy, White women. Health Psychology.

Ashar, Y. K., Andrews-Hanna, J. R., Dimidjian, S., & Wager, T. D. (2016). Towards a neuroscience of compassion: A brain systems-based model and research agenda. In J. D. Greene (Ed.), Positive Neuroscience (pp. 1–27). Oxford University Press.

Ashar, Y. K., Andrews-Hanna, J. R., Dimidjian, S., & Wager, T. D. (2017). Empathic Care and Distress: Predictive Brain Markers and Dissociable Brain Systems. Neuron, 1–11. http://doi.org/10.1016/j.neuron.2017.05.014

Ashar, Y. K., Andrews-Hanna, J. R., Yarkoni, T., Sills, J., Halifax, J., Dimidjian, S., & Wager, T. D. (2016). Effects of Compassion Meditation on a psychological model of charitable donation. Emotion.

Bloom, P. (2016). Against Empathy: The Case for Rational Compassion. Ecco.

Boot, W. R., Simons, D. J., Stothart, C., & Stutts, C. (2013). The Pervasive Problem With Placebos in Psychology: Why Active Control Groups Are Not Sufficient to Rule Out Placebo Effects. Perspectives on Psychological Science, 8(4), 445–454. http://doi.org/10.1177/1745691613491271

Cameron, D. (2017). Compassion Collapse, 1(July 2018), 1–21. http://doi.org/10.1093/oxfordhb/9780190464684.013.20

Cameron, D., Hutcherson, C., Ferguson, A., Andrew Scheffer, J., Hadjiandreou, E., & Inzlicht, M. (2019). Empathy is hard work: People choose to avoid empathy because of its cognitive costs. J Exp Psychol Gen. http://doi.org/10.31234/osf.io/jkc4n

Cutler, J., & Campbell-meiklejohn, D. (2019). A comparative fMRI meta-analysis of altruistic and strategic decisions to give. NeuroImage, 184(July 2018), 227–241. http://doi.org/10.1016/j.neuroimage.2018.09.009

Dahl, C. J., Lutz, A., & Davidson, R. J. (2015). Reconstructing and deconstructing the self: cognitive mechanisms in meditation practice. Trends in Cognitive Sciences, 19(9), 515–523. http://doi.org/10.1016/j.tics.2015.07.001

Dahl, C. J., Lutz, A., & Davidson, R. J. (2016). Cognitive Processes Are Central in Compassion Meditation. Trends in Cognitive Sciences, xx, 1–2. http://doi.org/10.1016/j.tics.2015.12.005

Engen, H. G., Bernhardt, B. C., Skottnik, L., Ricard, M., & Singer, T. (2018). Structural changes in socio-affective networks: Multi-modal MRI findings in long-term meditation practitioners. Neuropsychologia, 116(December 2016), 26–33. http://doi.org/10.1016/j.neuropsychologia.2017.08.024

Engen, H. G., & Singer, T. (2015a). Affect and motivation are critical in constructive meditation. Trends in Cognitive Sciences, xx, 1–2. http://doi.org/10.1016/j.tics.2015.11.004

Engen, H. G., & Singer, T. (2015b). Compassion-based emotion regulation up-regulates experienced positive affect and associated neural networks. Social Cognitive and Affective Neuroscience, 10(9), 1291–1301. http://doi.org/10.1093/scan/nsv008

Galante, J., Galante, I., Bekkers, M., & Gallacher, J. (2014). Effect of Kindness-Based Meditation on Health and Well-Being: A Systematic Review and Meta-Analysis. Journal of Consulting and Clinical Psychology.

Gilbert, P. (2014). The origins and nature of compassion focused therapy. British Journal of Clinical Psychology, 53(1), 6–41. http://doi.org/10.1111/bjc.12043

Glasser, M. F., Coalson, T. S., Robinson, E. C., Hacker, C. D., Harwell, J., Yacoub, E., … Van Essen, D. C. (2016). A multi-modal parcellation of human cerebral cortex. Nature, 1–11. http://doi.org/10.1038/nature18933

Goetz, J. L., Keltner, D., & Simon-Thomas, E. (2010). Compassion: an evolutionary analysis and empirical review. Psychological Bulletin, 136(3), 351–74. http://doi.org/10.1037/a0018807

Haber, S. N., & Knutson, B. (2010). The Reward Circuit: Linking Primate Anatomy and Human Imaging. Neuropsychopharmacology, 35(1), 4–26. http://doi.org/10.1038/npp.2009.129

Halifax, J. (2012). A heuristic model of enactive compassion. Current Opinion in Supportive and Palliative Care, 6(2), 228–35. http://doi.org/10.1097/SPC.0b013e3283530fbe

Kanske, P., Bockler, A., Trautwein, F. M., Lesemann, F. H. P., & Singer, T. (2016). Are strong empathizers better mentalizers? Evidence for independence and interaction between the routes of social cognition. Social Cognitive and Affective Neuroscience, 11(9), 1383–1392. http://doi.org/10.1093/scan/nsw052

Kaptchuk, T. J., Kelley, J. M., Conboy, L. A., Davis, R. B., Kerr, C. E., Jacobson, E. E., … Lembo, A. J. (2008). Components of placebo effect: randomised controlled trial in patients with irritable bowel syndrome. BMJ, 336(7651), 999–1003. http://doi.org/10.1136/bmj.39524.439618.25

Klimecki, O. M., Leiberg, S., Lamm, C., & Singer, T. (2012). Functional Neural Plasticity and Associated Changes in Positive Affect After Compassion Training. Cerebral Cortex (New York, N.Y.: 1991), 23, 1–10. http://doi.org/10.1093/cercor/bhs142

Klimecki, O. M., Leiberg, S., Ricard, M., & Singer, T. (2014). Differential pattern of functional brain plasticity after compassion and empathy training. Social Cognitive and Affective Neuroscience, 9(6), 873–9. http://doi.org/10.1093/scan/nst060

Klimecki, O. M., & Singer, T. (2011). Empathic Distress Fatigue Rather Than Compassion Fatigue? Integrating Findings from Empathy Research in Psychology and Social Neuroscience. In B. Oakley, A. Knafo, G. Madhavan, & D. S. Wilson (Eds.), Pathological altruism (pp. 368–383). New York: Oxford University Press.

Kok, B. E., & Singer, T. (2017). Phenomenological Fingerprints of Four Meditations: Differential State Changes in Affect, Mind-Wandering, Meta-Cognition, and Interoception Before and After Daily Practice Across 9 Months of Training. Mindfulness, 8(1), 218–231. http://doi.org/10.1007/s12671-016-0594-9

Kreplin, U., Farias, M., & Brazil, I. A. (2018). The limited prosocial effects of meditation: A systematic review and meta-analysis. Scientific Reports, 8(1), 1–10. http://doi.org/10.1038/s41598-018-20299-z

MacCoon, D. G., Imel, Z. E., Rosenkranz, M. A., Sheftel, J. G., Weng, H. Y., Sullivan, J. C., … Lutz, A. (2012). The validation of an active control intervention for Mindfulness Based Stress Reduction (MBSR). Behaviour Research and Therapy, 50(1), 3–12. http://doi.org/10.1016/j.brat.2011.10.011

Mascaro, J. S., Kelley, S., Darcher, A., Negi, L. T., Worthman, C., Miller, A., & Raison, C. (2018). Meditation buffers medical student compassion from the deleterious effects of depression. Journal of Positive Psychology, 13(2), 133–142. http://doi.org/10.1080/17439760.2016.1233348

Mascaro, J. S., Rilling, J. K., Tenzin Negi, L., & Raison, C. L. (2013a). Compassion meditation enhances empathic accuracy and related neural activity. Social Cognitive and Affective Neuroscience, 8(1), 48–55. http://doi.org/10.1093/scan/nss095

Mascaro, J. S., Rilling, J. K., Tenzin Negi, L., & Raison, C. L. (2013b). Compassion meditation enhances empathic accuracy and related neural activity. Social Cognitive and Affective Neuroscience, 8(1), 48–55. http://doi.org/10.1093/scan/nss095

Maslach, C., & Leiter, M. P. (2016). Understanding the burnout experience: Recent research and its implications for psychiatry. World Psychiatry, 15(2), 103–111. http://doi.org/10.1002/wps.20311

Nieuwenhuis, S., Forstmann, B. U., & Wagenmakers, E. (2011). Erroneous analyses of interactions in neuroscience: a problem of significance. Nature Neuroscience, 14(9), 1105–1109. http://doi.org/10.1038/nn.2886

Pace, T. W. W., Negi, L. T., Adame, D. D., Cole, S. P., Sivilli, T. I., Brown, T. D., … Raison, C. L. (2009). Effect of compassion meditation on neuroendocrine, innate immune and behavioral responses to psychosocial stress. Psychoneuroendocrinology, 34(1), 87–98. http://doi.org/10.1016/j.psyneuen.2008.08.011

Pauli, W. M., Nili, A. N., & Tyszka, J. M. (2018). A high-resolution probabilistic in vivo atlas of human subcortical brain nuclei. Scientific Data, 5, 180063.

Rakel, D. D., Barrett, B., Zhang, Z., Hoeft, T., Chewning, B., Marchand, L., … Scheder, J. (2011). Perception of empathy in the therapeutic encounter: effects on the common cold. Patient Education and Counseling, 85(3), 390–7. http://doi.org/10.1016/j.pec.2011.01.009

Segal, Z. V, Bieling, P., Young, T., MacQueen, G., Cooke, R., Martin, L., … Levitan, R. D. (2010). Antidepressant monotherapy vs sequential pharmacotherapy and mindfulness-based cognitive therapy, or placebo, for relapse prophylaxis in recurrent depression. Archives of General Psychiatry, 67(12), 1256–64. http://doi.org/10.1001/archgenpsychiatry.2010.168

Shanafelt, T., Goh, J., & Sinsky, C. (2017). The business case for investing in physician well-being. JAMA Internal Medicine, 177(12), 1826–1832. http://doi.org/10.1001/jamainternmed.2017.4340

Singer, T., & Klimecki, O. M. (2014). Empathy and compassion. Current Biology: CB, 24(18), R875–8. http://doi.org/10.1016/j.cub.2014.06.054

Valk, S. L., Bernhardt, B. C., Trautwein, F. M., Böckler, A., Kanske, P., Guizard, N., … Singer, T. (2017). Structural plasticity of the social brain: Differential change after socio-affective and cognitive mental training. Science Advances, 3(10), 1–12. http://doi.org/10.1126/sciadv.1700489

van Berkhout, E. T., & Malouff, J. M. (2015). The Efficacy of Empathy Training: A Meta-Analysis of Randomized Controlled Trials. Journal of Counseling Psychology.

Wager, T. D., Keller, M. C., Lacey, S. C., & Jonides, J. (2005). Increased sensitivity in neuroimaging analyses using robust regression. NeuroImage, 26(1), 99–113. http://doi.org/10.1016/j.neuroimage.2005.01.011

Watson, K. K., & Platt, M. L. (2012). Social Signals in Primate Orbitofrontal Cortex. Current Biology, 22(23), 2268–2273. http://doi.org/10.1016/j.cub.2012.10.016

Weng, H. Y., Fox, A. S., Shackman, A. J., Stodola, D. E., Caldwell, J. Z. K., Olson, M. C., … Davidson, R. J. (2013). Compassion Training Alters Altruism and Neural Responses to Suffering. Psychological Science. http://doi.org/10.1177/0956797612469537

Weng, H. Y., Schuyler, B., & Davidson, R. J. (2017). The Impact of Compassion Meditation Training on the Brain and Prosocial Behavior. In The Oxford Handbook of Compassion Science.

Williams, J. M. G., Crane, C., Barnhofer, T., Brennan, K., Duggan, D. S., Fennell, M. J. V, … Russell, I. T. (2014). Mindfulness-based cognitive therapy for preventing relapse in recurrent depression: a randomized dismantling trial. Journal of Consulting and Clinical Psychology, 82(2), 275–86. http://doi.org/10.1037/a0035036

Woo, C. W., Krishnan, A., & Wager, T. D. (2014). Cluster-extent based thresholding in fMRI analyses: Pitfalls and recommendations. NeuroImage, 91, 412–419. http://doi.org/10.1016/j.neuroimage.2013.12.058

Zajonc, R. B. (2001). Mere Exposure: A Gateway to the Subliminal. Current Directions in Psychological Science, 10(6), 224–228. http://doi.org/10.1111/1467-8721.00154

Zaki, J. (2014). Empathy: A Motivated Account. Psychological Bulletin, 140(6), 1608–1647.

Zeidan, F., Emerson, N. M., Farris, S. R., Ray, J. N., Jung, Y., McHaffie, J. G., & Coghill, R. C. (2015). Mindfulness Meditation-Based Pain Relief Employs Different Neural Mechanisms Than Placebo and Sham Mindfulness Meditation-Induced Analgesia. Journal of Neuroscience, 35(46), 15307–15325. http://doi.org/10.1523/JNEUROSCI.2542-15.2015

Zeidan, F., Johnson, S. K., Gordon, N. S., & Goolkasian, P. (2010). Effects of brief and sham mindfulness meditation on mood and cardiovascular variables. Journal of Alternative and Complementary Medicine (New York, N.Y.), 16(8), 867–873. http://doi.org/10.1089/acm.2009.0321

