## Supplementary materials for "Effects of compassion training on brain responses to suffering others"

Supp. Table 1

| Region | Volume (mm^3^) | *x* | *y* | *z* | max/min *Z* |
| --- | --- | --- | --- | --- | --- |
| **CM vs. Familiarity** |  |  |  |  |  |
| *CM > Familiarity* |  |  |  |  |  |
| L STS | 336 | -56 | -6 | -14 | 3.67 |
| R mOFC | 288 | 8 | 18 | -14 | 3.67 |
| L parahippocampal cortex | 528 | -18 | -38 | -16 | 3.90 |
| *Familiarity > CM* |  |  |  |  |  |
| L PFC | 704 | -44 | 38 | 36 | 3.65 |
| **CM vs. OxyPla** |  |  |  |  |  |
| *CM > OxyPla* |  |  |  |  |  |
| R mOFC | 720 | 10 | 20 | -16 | 3.90 |
| R V1 | 1904 | 6 | -66 | 18 | 3.86 |
| L V3 | 464 | -10 | -96 | 34 | 3.62 |
| *OxyPla > CM* |  |  |  |  |  |
| L V1 | 512 | -18 | -96 | -10 | 3.90 |
| **OxyPla vs. Familiarity** |  |  |  |  |  |
| *OxyPla > Familiarity* |  |  |  |  |  |
| L midtemporal gyrus | 424 | -44 | -10 | -16 | 3.87 |
| *Familiarity > OxyPla* |  |  |  |  |  |
| R V1 | 1464 | 8 | -68 | 18 | -3.91 |

Note. Group differences in pre-to-post intervention changes in brain responses to stories of suffering, for the CM vs. Familiarity and CM vs. placebo oxytocin comparisons (Figure 1A).

Supp. Table 2

| Region | Volume (mm^3^) | *x* | *y* | *z* | max/min *Z* |
| --- | --- | --- | --- | --- | --- |
| **CM** |  |  |  |  |  |
| *Increases* |  |  |  |  |  |
| L lateral OFC | 336 | -20 | 18 | -18 | 4.01 |
| L mPFC | 472 | -12 | 50 | 8 | 4.47 |
| R cerebellum | 720 | 16 | -34 | -32 | 4.79 |
| Midbrain | 640 | 6 | -26 | 2 | 4.78 |
| *Decreases* |  |  |  |  |  |
| R STS | 1200 | 58 | -22 | 12 | -4.38 |
| R mid-temporal gyrus | 1184 | 62 | -36 | 4 | -5.05 |
| R occipital | 344 | 40 | -80 | -14 | -4.58 |
| V4 | 1864 | -32 | -94 | 12 | -5.00 |
| V4 | 776 | 36 | -92 | 10 | -3.70 |
| **OxyPla** |  |  |  |  |  |
| *Increases* |  |  |  |  |  |
| - |  |  |  |  |  |
| *Decreases* |  |  |  |  |  |
| L V3 | 312 | -4 | -96 | 32 | -3.755 |
| R Frontal eye field | 352 | 38 | -4 | 44 | -5.6535 |
| R V3 | 768 | 4 | -94 | 22 | -3.7862 |
| L midfrontal gyrus | 416 | -38 | 8 | 64 | -3.7456 |
| **Familiarity** |  |  |  |  |  |
| *Increases* |  |  |  |  |  |
| L midfrontal gyrus | 736 | -42 | 40 | 36 | 3.86 |
| *Decreases* |  |  |  |  |  |
| R inf. long. fasciculus | 304 | 48 | -22 | -16 | -5.0978 |
| R hippocampus | 464 | 30 | -18 | -14 | -3.6779 |
| R STS | 328 | 62 | -38 | 0 | -6.0503 |
| L temporal pole | 272 | -42 | 6 | -20 | -3.7768 |
| L hypothalamus | 1776 | -4 | -6 | -16 | -5.1774 |
| L hypothalamus | 1144 | -4 | -2 | -4 | -6.1546 |
| L mOFC | 352 | -8 | 8 | -14 | -4.5446 |
| R insula | 368 | 38 | 34 | 2 | -4.904 |
| Inferior temporal gyrus | 464 | -46 | -4 | -36 | -4.8132 |
| L amygdala | 384 | -16 | -2 | -16 | -5.231 |
| L external globus pallidus | 2048 | -20 | 6 | -8 | -4.464 |
| R external globus pallidus | 1192 | 16 | 8 | -2 | -4.4978 |
| Putamen | 1144 | -30 | -14 | -10 | -4.3297 |
| Putamen | 504 | 30 | -18 | 0 | -4.1317 |

Note. Absolute pre-to-post intervention changes in brain responses to stories of suffering, for each condition separately (Figure 1B). See max/min Z to indicate increase vs. decrease.

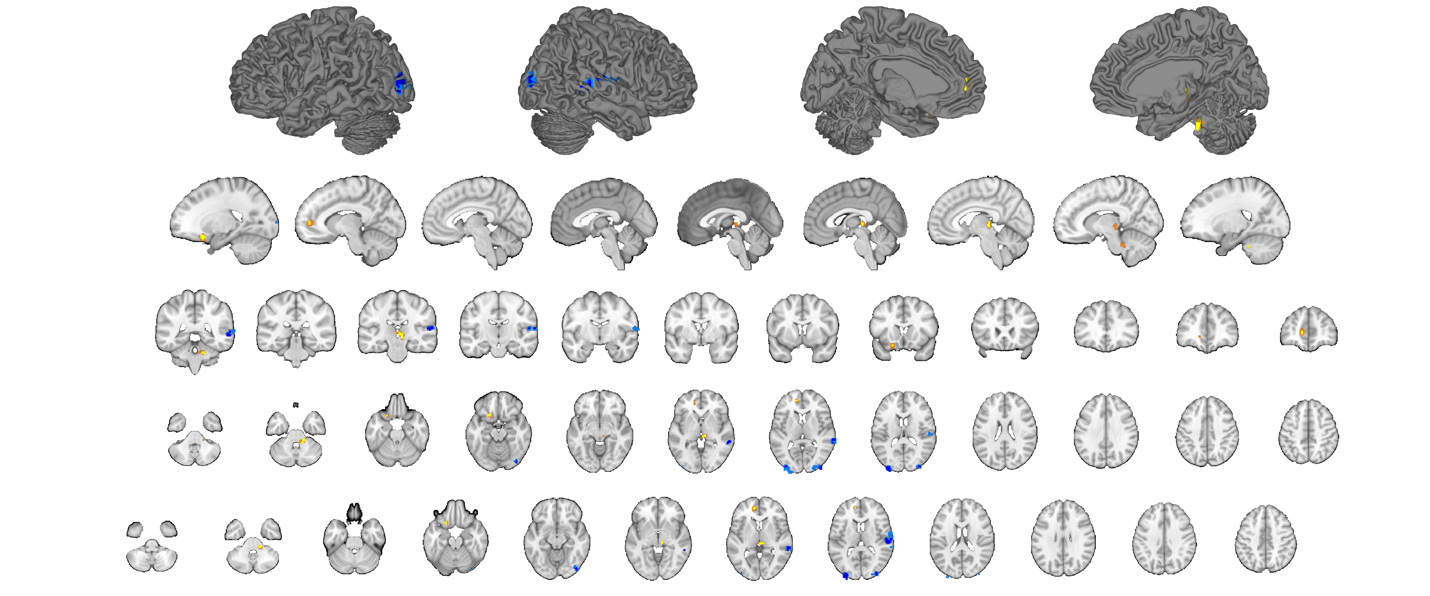

Figure S1. Whole brain pre-to-post-intervention changes in the CM condition. Voxels meeting a threshold of *p* < .001 uncorrected are shown in yellow and dark blue, respectively. Adjacent voxels meeting a threshold of *p* < .005 are shown in orange/light blue, for visualization purposes.

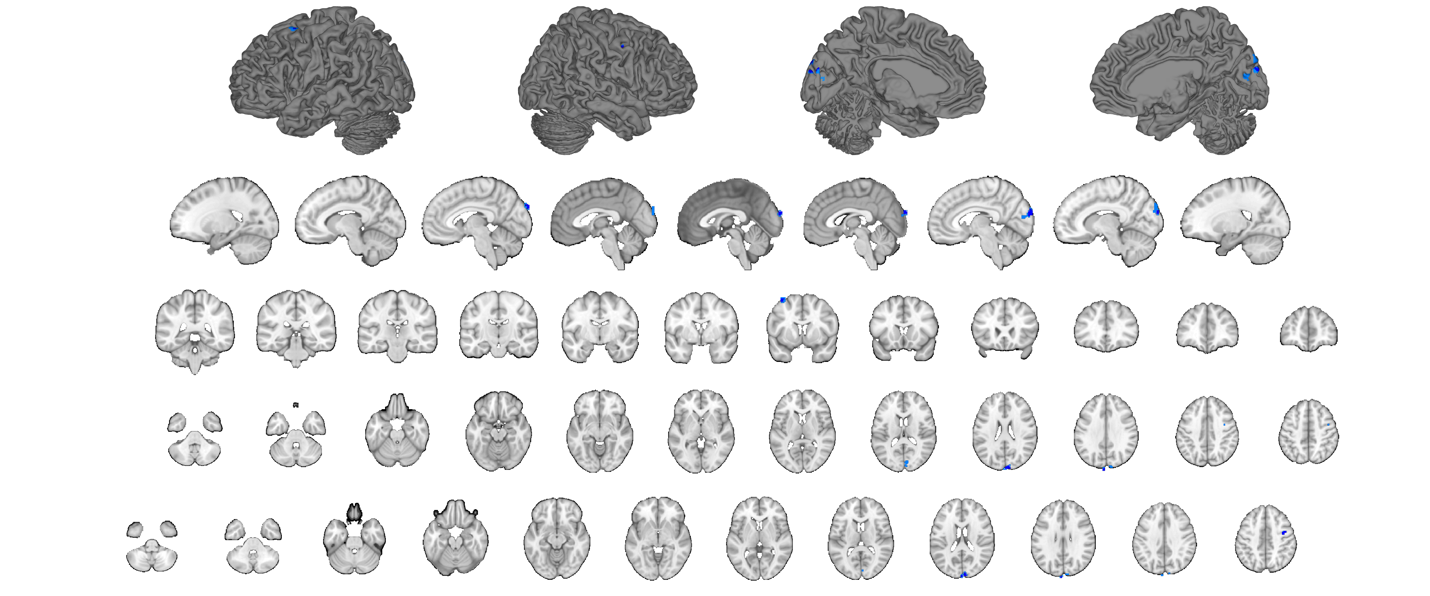

Figure S2. Whole brain pre-to-post-intervention changes in the OxyPla condition. Voxels meeting a threshold of *p* < .001 uncorrected are shown in yellow and dark blue, respectively. Adjacent voxels meeting a threshold of *p* < .005 are shown in orange/light blue, for visualization purposes.

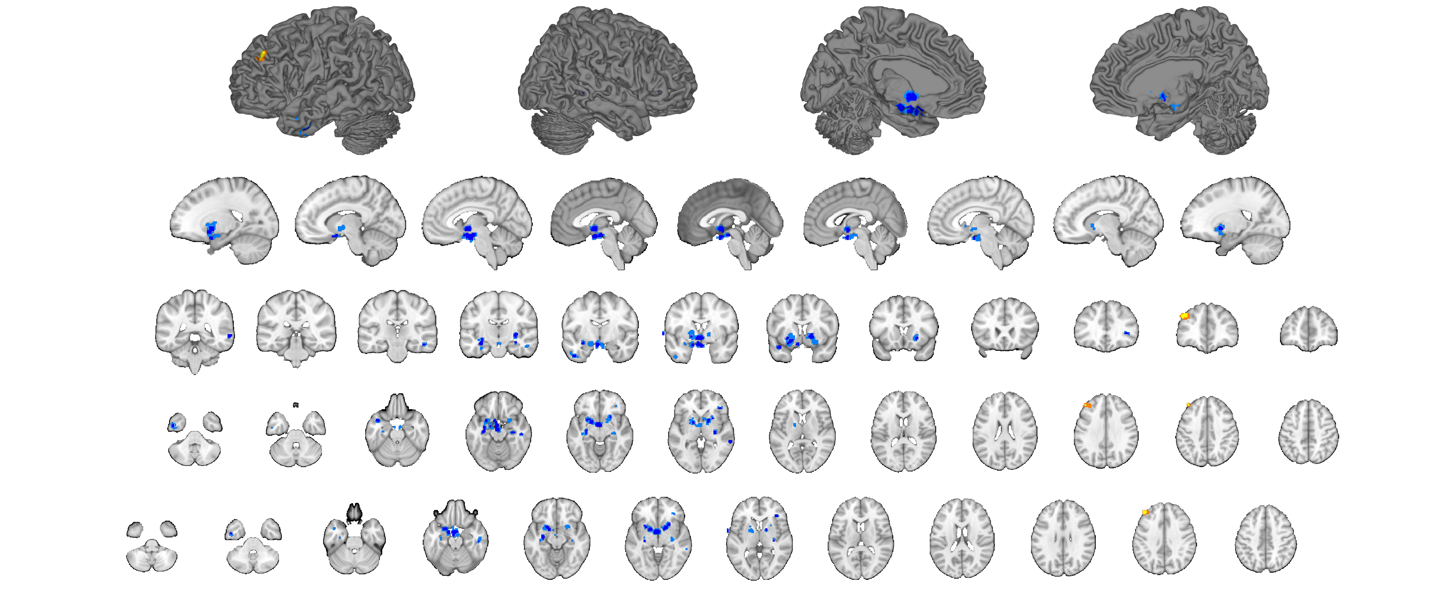

Figure S3. Whole brain pre-to-post-intervention changes in the Familiarity condition. Voxels meeting a threshold of *p* < .001 uncorrected are shown in yellow and dark blue, respectively. Adjacent voxels meeting a threshold of *p* < .005 are shown in orange/light blue, for visualization purposes.

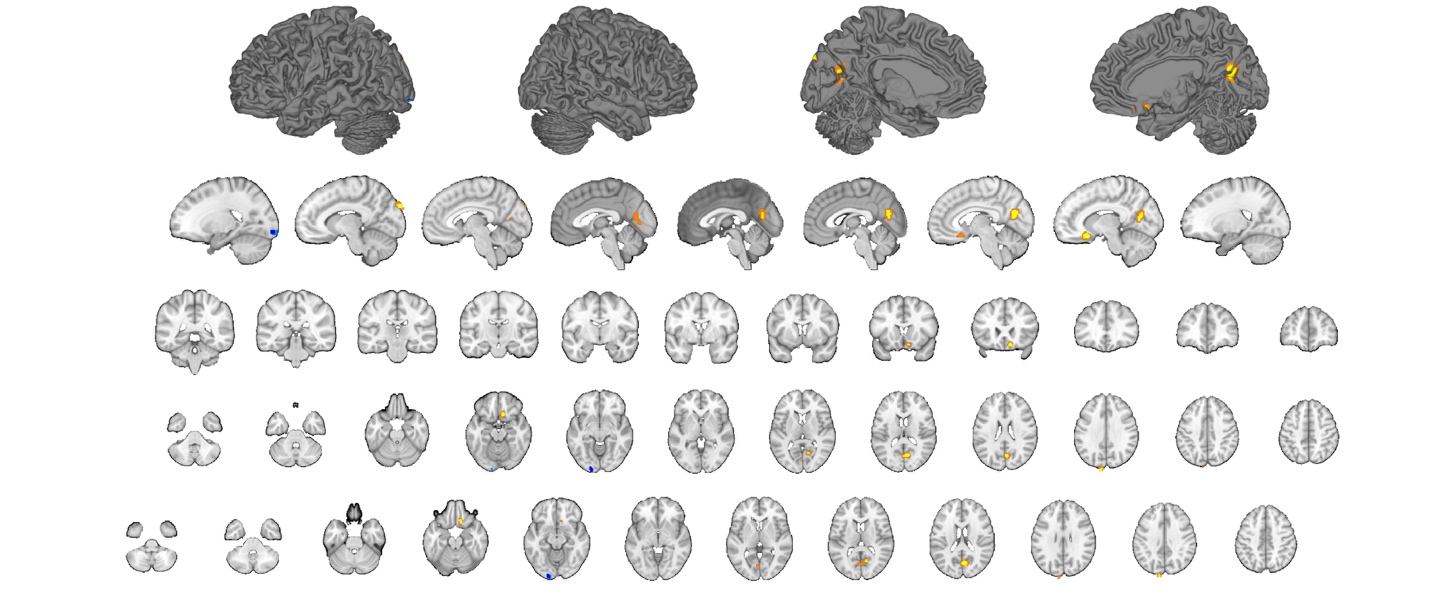

Figure S4. Whole brain results for CM vs. OxyPla differences in pre-to-post-intervention changes. Voxels meeting a threshold of *p* < .001 uncorrected are shown in yellow and dark blue, respectively. Adjacent voxels meeting a threshold of *p* < .005 are shown in orange/light blue, for visualization purposes.

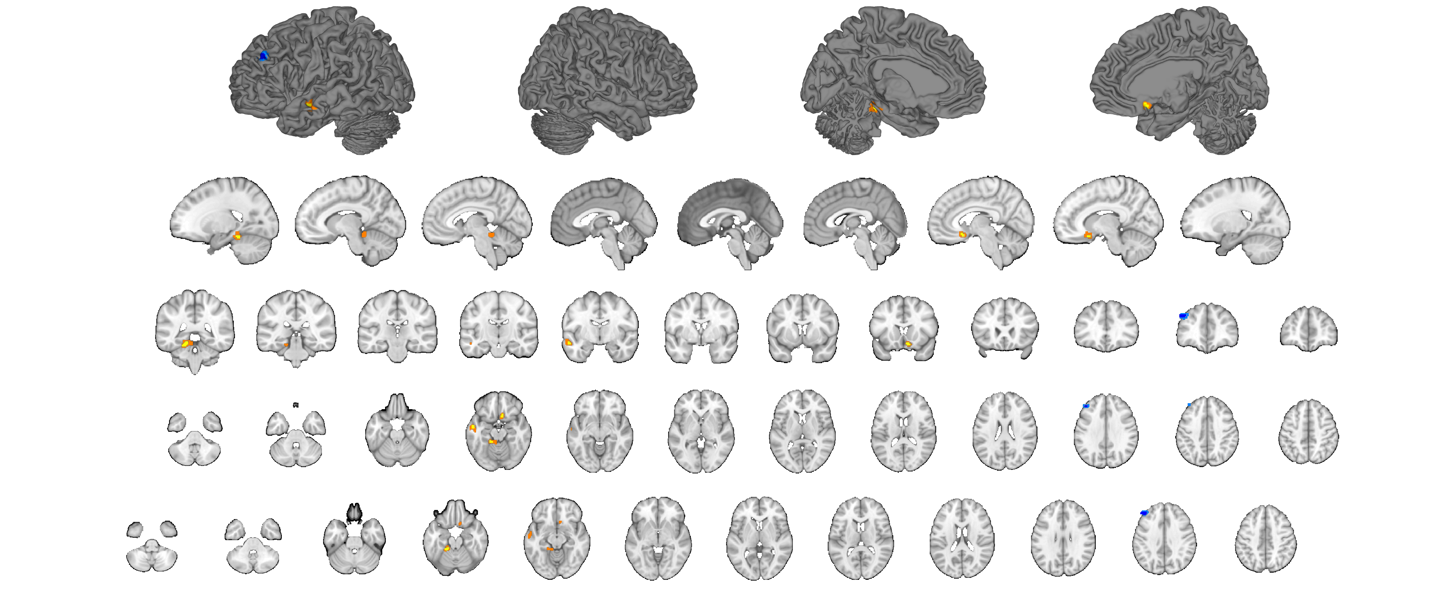

Figure S5. Whole brain results for CM vs. Familiarity differences in pre-to-post-intervention changes. Voxels meeting a threshold of *p* < .001 uncorrected are shown in yellow and dark blue, respectively. Adjacent voxels meeting a threshold of *p* < .005 are shown in orange/light blue, for visualization purposes.

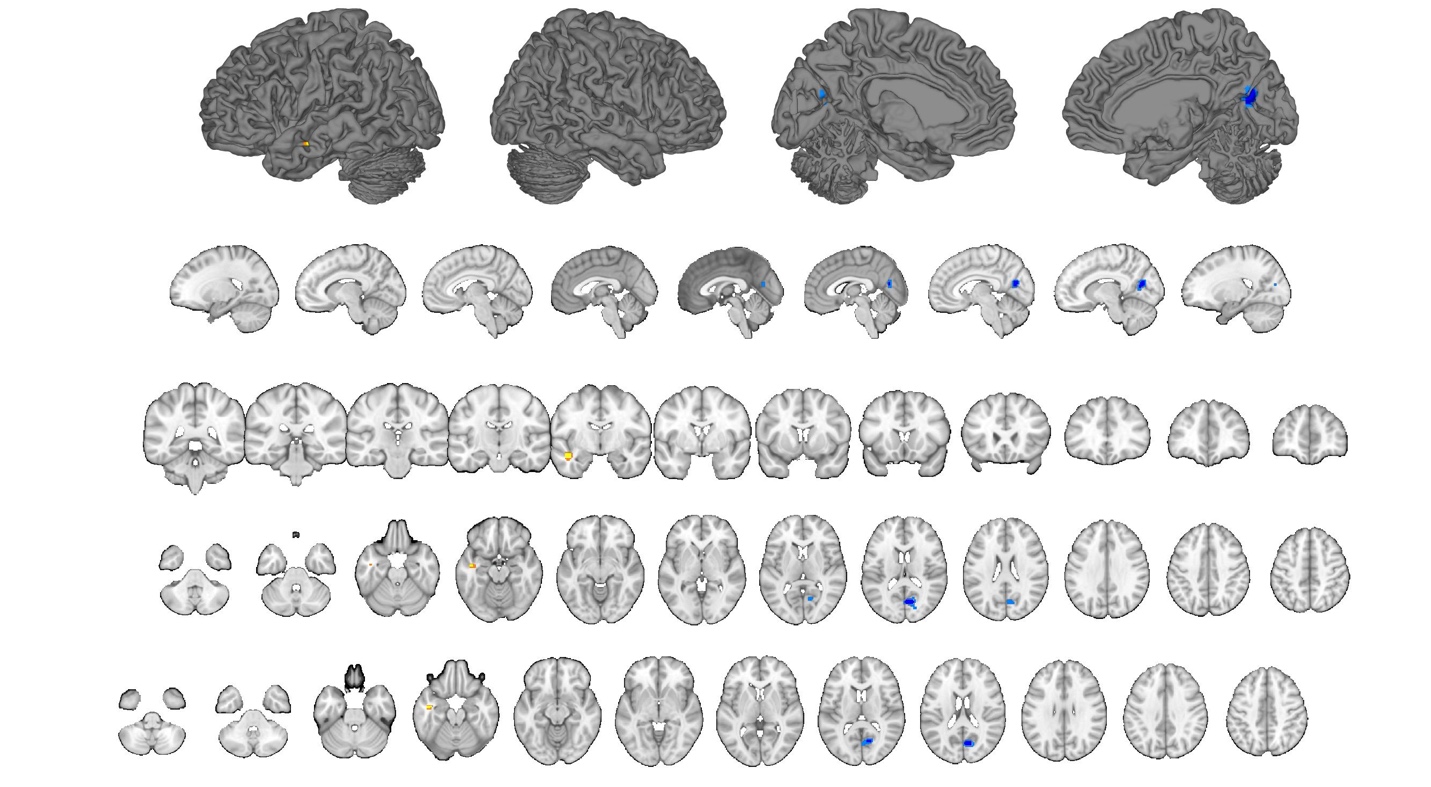

Figure S6. Whole brain results for OxyPla vs. Familiarity differences in pre-to-post-intervention changes. Voxels meeting a threshold of *p* < .001 uncorrected are shown in yellow and dark blue, respectively. Adjacent voxels meeting a threshold of *p* < .005 are shown in orange/light blue, for visualization purposes.
